# Modeling multiple phenotypes in wheat using data-driven genomic exploratory factor analysis and Bayesian network learning

**DOI:** 10.1101/2020.09.03.282335

**Authors:** Mehdi Momen, Madhav Bhatta, Waseem Hussain, Haipeng Yu, Gota Morota

**Affiliations:** Department of Animal and Poultry Sciences, Virginia Polytechnic Institute and State University, Blacksburg, VA, USA 24061; Department of Agronomy, University of Wisconsin-Madison, Madison, WI 53706, USA; International Rice Research Institute, Los Banos, Philippines

**Keywords:** Bayesian network, confirmatory factor analysis, exploratory factor analysis, multi-trait, wheat

## Abstract

Inferring trait networks from a large volume of genetically correlated diverse phenotypes such as yield, architecture, and disease resistance can provide information on the manner in which complex phenotypes are interrelated. However, studies on statistical methods tailored to multi-dimensional phenotypes are limited, whereas numerous methods are available for evaluating the massive number of genetic markers. Factor analysis operates at the level of latent variables predicted to generate observed responses. The objectives of this study were to illustrate the manner in which data-driven exploratory factor analysis can map observed phenotypes into a smaller number of latent variables and infer a genomic latent factor network using 45 agro-morphological, disease, and grain mineral phenotypes measured in synthetic hexaploid wheat lines (*Triticum Aestivum L.*). In total, eight latent factors including grain yield, architecture, flag leaf-related traits, grain minerals, yellow rust, two types of stem rust, and leaf rust were identified as common sources of the observed phenotypes. The genetic component of the factor scores for each latent variable was fed into a Bayesian network to obtain a trait structure reflecting the genetic interdependency among traits. Three directed paths were consistently identified by two Bayesian network algorithms. Flag leaf-related traits influenced leaf rust, and yellow rust and stem rust influenced grain yield. Additional paths that were identified included flag leaf-related traits to minerals and minerals to architecture. This study shows that data-driven exploratory factor analysis can reveal smaller dimensional common latent phenotypes that are likely to give rise to numerous observed field phenotypes without relying on prior biological knowledge. The inferred genomic latent factor structure from the Bayesian network provides insights for plant breeding to simultaneously improve multiple traits, as an intervention on one trait will affect the values of focal phenotypes in an interrelated complex trait system.

## Background

With the development of high-throughput phenotyping technologies, phenomics has been generating plant measurements at a greater level of resolution and dimensionality (Araus and Cairns, 2014; Watanabe et al., 2017). Integrating these diverse and heterogeneous data to improve the biological understanding of plant systems and interpret the underlying inter-relationships among phenotypes remains challenging (Morota et al., 2019). One approach is to model each measurement as a different trait using a multi-trait model (Henderson and Quaas, 1976). However, in a high-dimensional specification, where the number of traits measured per genotype can reach hundreds or thousands, this approach leads to dramatic increases in the computational burden or difficulties in interpreting the results. Recently, Yu et al. (2019) showed that factor analysis can be used to reduce the dimension of response variables by assuming latent factors that give rise to observed phenotypes in rice. They used confirmatory factor analysis (CFA), which requires knowledge of the phenotype-factor category before data analysis. However, reliable phenotype-factor patterns are not always known in advance. Alternatively, exploratory factor analysis (EFA) can be used to perform latent variable analysis by estimating patterns from data when a latent structure cannot be determined a priori. EFA identifies underlying latent factors to represent observed measurements, which is useful when the exact number and meaning of latent factors are unknown (Jöreskog, 1967; Hoyle and Duvall, 2004).

The first objective of this study was to illustrate the utility of EFA for revealing the underlying genomic latent structure of agronomic or agro-morphological phenotypes for synthetic hexaploid wheat lines (*T. aestivum L*). Grain yield in wheat is influenced by several agro-morphological traits. However, successfully incorporating yield-promoting agro-morphological traits in breeding programs to improve genetic gains requires detailed knowledge of the interrelationships between and among traits. The second objective was to determine a trait network structure among the genomic latent factors using a Bayesian network. This is an essential task because breeding programs often aim to improve multiple correlated traits concurrently. Knowledge of directed trait networks accounting for the genetic interdependency among traits can improve the understanding of the manner in which the selection of one phenotype may increase or decrease the observation of another phenotype, providing additional insight beyond associations (Valente et al., 2015). The current study demonstrates the advantages of the joint application of factor analysis and Bayesian network as a data-driven approach to discover interrelationships between a set of many correlated traits in wheat.

## Materials and Methods

### Plant materials

A diversity panel of *n* = 123 synthetic hexaploid wheat lines, derived from an interspecific cross between wild accessions of goat grass (*Aegilops tauschii L.*) and diverse accessions of cultivated durum wheat (*Triticum turgidum L.*), was used in this study. These plant materials were shared by the International Winter Wheat Improvement Program in Turkey and are available at http://www.iwwip.org. Pedigree information and other details on these lines were reported previously (Bhatta et al., 2018a,c,d). Briefly, the lines originated from two breeding programs. The first group of synthetics comprises 14 lines developed by Kyoto University, Japan, from 1 Langdon durum parent crossed with 14 different accessions of *Ae. tauschii*. The second group consists of 109 lines developed by the International Maize and Wheat Improvement Center from crosses between 6 winter durum wheats and 11 different *Ae. tauschii* accessions. The synthetic lines used in this study are unique; they were recently developed (F8–F9 generations) and tested for multiple traits for use in a breeding program.

### Phenotypic and genotypic data

We analyzed 16 agronomic-, 16 grain mineral-, and 13 wheat rust-related phenotypes in the current study. Agronomic traits including grain yield (GY), harvest index (HI), biomass weight (BMWT), grain volume weight (GVWT), flag leaf length (FLL), flag leaf width (FLW), flag leaf area (FLA), rachis break (RB), sterile spikelet (SP), spike length (SL), seeds per spike (SPS), spikelet number (SN), fertile spikelet (FS), spike weight (SW), grain weight per spike (GPS), and spike harvest index (SHI) were measured using previously described standard procedures (Bhatta et al., 2018a; Morgounov et al., 2018; Hussain et al., 2017). Grain minerals including arsenic (As), calcium (Ca), cadmium (Cd), cobalt (Co), copper (Cu), iron (Fe), potassium (K), lithium (Li), magnesium (Mg), manganese (Mn), molybdenum (Mo), nickel (Ni), phosphorous (P), sulfur (S), titanium (Ti), and zinc (Zn) were measured via inductively-coupled plasma mass spectrometry (ICP-MS, Agilent 7500cx, Agilent Technologies, Santa Clara, CA, USA) at the University of Nebraska Redox Biology Center, Proteomics and Metabolomics Core (Guttieri et al., 2015; Bhatta et al., 2018a). The wheat rust (leaf stem and yellow rusts) disease severity, coefficient of infection, and infection type were tested under field conditions as previously described (Peterson et al., 1948; Morgounov et al., 2018; Bhatta et al., 2018d). Wheat rust traits collected from several locations in Turkey and one location in Kenya included the leaf rust coefficient of infection (LRCI), leaf rust infection type (LRIT), leaf rust severity (LRS), stem rust coefficient of infection at Haymana (SRCIH), stem rust infection type at Haymana (SRITH), stem rust severity at Haymana (SRSH), stem rust coefficient of infection at Kastamonu (SRCIK), stem rust infection type at Kastamonu (SRITK), stem rust severity at Kastamonu (SRSK), yellow rust coefficient of infection at Haymana (YRCIH), yellow rust infection type at Haymana (YRIH), yellow rust severity at Haymana (YRSH), and yellow rust severity at Kastamonu (YRSK). All lines were genotyped with the genotyping by sequencing technology (Bhatta et al., 2018c). After setting a minor allele frequency threshold of 0.05, 35,648 markers remained for analysis.

### Experimental design and analysis

The experiments were conducted across several locations in Turkey and one location in Kenya in 2017. The experimental design was an alpha lattice design with two replications (Barreto et al., 1996). A linear mixed model coupled with restricted maximum likelihood implemented in the PROC MIXED procedure in SAS 9.4 (SAS Institute, Inc., Cary, NC, USA) was used to obtain the adjusted means for each trait from the following model (Bhatta et al., 2018b).

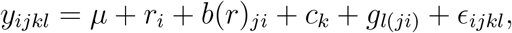

where *y_ijk_* is the trait of interest; *μ* is the overall mean; *r_i_* is the effect of *i*th replication; *b*(*r*)_*ji*_ is the effect of the *j*th block within the *i*th replication; *c_k_* is the *k*th check; *g_lji_* (new variable, where check is coded as 0 and entry is coded as 1, and the genotype is considered a new variable × entry) is the effect of the *l*th genotype within the *j*th incomplete block of the *i*th replication; and *ϵ_ijkl_* is the residual.

### Exploratory factor analysis

Exploratory factor analysis can reveal the latent structure among phenotypes when no hypotheses about the nature of the underlying factor can be assumed *a priori*. This section closely follows the work of Yu et al. (2020). The aforementioned *t* = 45 phenotypes were analyzed using EFA by fitting

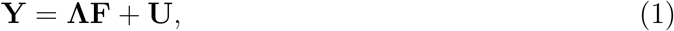

where **Y** is the *t* × *n* phenotypic matrix; **Λ** is the *t* × *q* matrix of factor loading indicating the relation between phenotypes and latent common factors; **F** is the *q* × *n* matrix of latent factor scores; and **U** is the *t* × *n* vector of unique effects that is not explained by *q* underlying common factors. The variance-covariance matrix of **Y** is

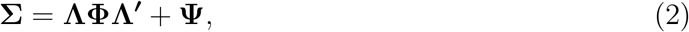

where **Σ** is the *t* × *t* variance-covariance matrix of phenotypes, **Φ** is the variance of factor scores, and **Ψ** is a *t* × *t* diagonal matrix of unique variance. The elements of **Λ**, **Φ**, and **Ψ** are parameters of the model to be estimated from the data. We assumed **Φ** = **I** yielding factors each with unit variance (Jöreskog, 1967; Anderson, 2003). With the assumption of 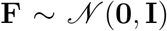, parameters **Λ** and **Ψ** were estimated by maximizing the log-likelihood of 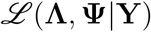 using the R package psych (Revelle, 2018) along with a varimax rotation (Kaiser, 1958). A threshold of λ > |0.3| was first applied to screen out factor loading values. Then each phenotype was assigned to only one of the factors based on its largest loading.

Parallel analysis was performed to estimate the optimum number of factors from data in EFA (Horn, 1965; Hayton et al., 2004). This is conducted by generating simulated data from the observed data. Next, the eigenvalues were extracted until the observed data had a smaller eigenvalue than the simulated data. The number of eigenvalues was used as the number of optimum factors.

The factor ability of the data set was also assessed by estimating the Kaiser-Meyer-Olkin measure of sampling adequacy (Cerny and Kaiser, 1977). This criterion measures the adequacy of the dataset for factor analysis by investigating the correlation and partial correlation matrices of the phenotypes. The measure of sampling adequacy ranges between 0 to 1, and values closer to 1 are preferred. When the measure of sampling adequacy is less than 0.5, the dataset is not recommended for factor analysis (Cerny and Kaiser, 1977).

### Confirmatory factor analysis

Once the phenotype-factor pattern was established by EFA, Bayesian CFA was used to obtain factor scores. Although EFA and CFA are similar, there are also clear differences. In general, EFA is used to find a latent structure in data, whereas CFA requires the phenotype-latent variable category to be known before analysis and is often used to estimate factor scores based on the structure from EFA. The differences between EFA and CFA are shown in Figure 1. In a Bayesian setting, all unknowns in equations (1) and (2) were assigned priors. The assignment of priors was performed according to Yu et al. (2019, 2020) using the default priors in the blavaan R package (Merkle and Rosseel, 2018). A Gaussian distribution with a mean of zero and variance of 100 was assigned to the factor loading term. The variance-covariance matrix of the latent factors followed an inverse Wishart distribution with a scale matrix of an 8 × 8 identity matrix and degree of freedom of 8. Each error variance followed an inverse Gamma distribution with a shape parameter of 1 and scale parameter of 0.5. The factor scores of latent variables (**F**) were sampled from the conditional distribution of *p*(**F**|**Λ**, **Φ**, **Ψ**, **Y**) (Lee and Song, 2012) using a data augmentation technique (Tanner and Wong, 1987). The posterior mean of **F** was considered a new phenotype in subsequent analysis. Convergence was diagnosed by the potential scale reduction factor (PSRF) (Gelman et al., 1992; Brown, 2014). This criterion utilizes at least two Markov chains, which are considered to be mixed to a stationary status if the ratio of between the chain variance to within the chain variance is close to 1. In total, two chains, each consisting of 5,000 Markov chain Monte Carlo samples after 2,000 burn-in samples, were collected to derive the posterior means.

**Figure 1:**
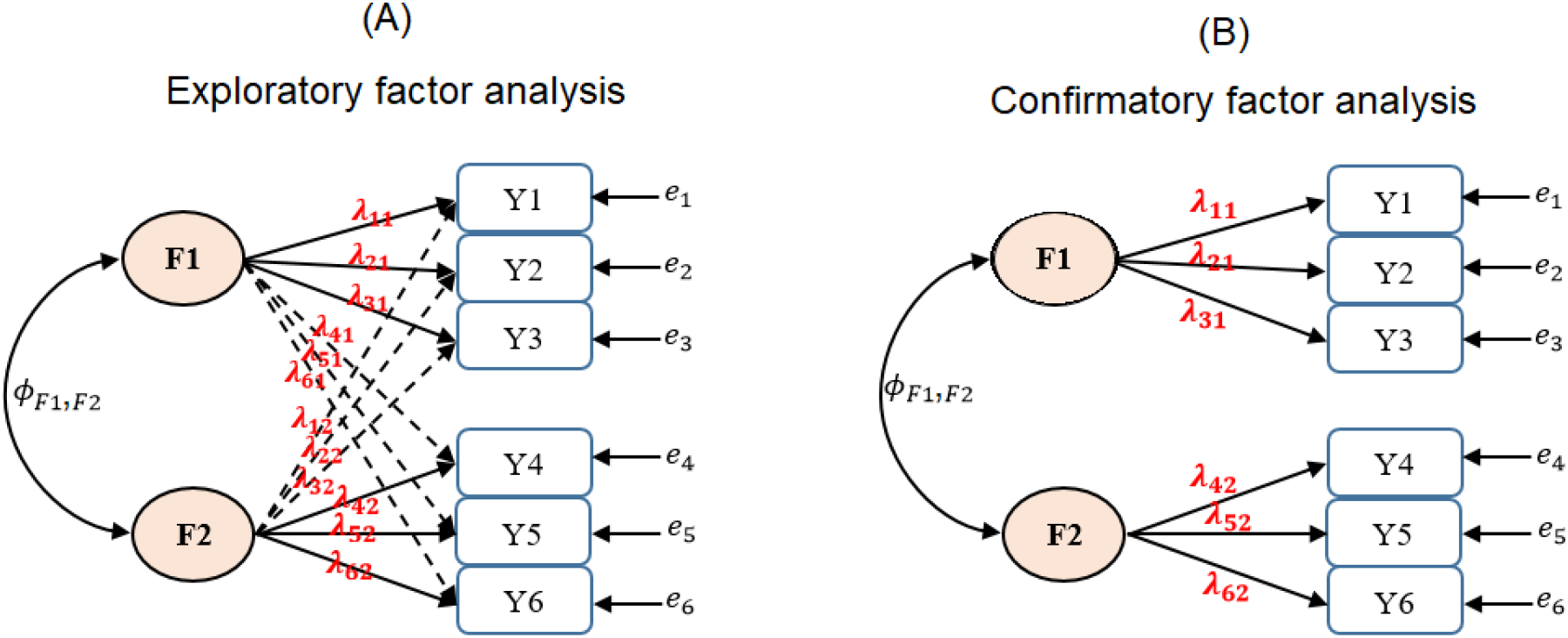
A graphical representation of exploratory factor analysis (panel A) and confirmatory factor analysis (panel B) assuming that there are hypothetical six observed phenotypes (Y1, Y2, ⋯, Y6) and two unobserved latent factors (F1 and F2). The double headed arrow is the covariance between the two latent factors (Φ_*F*1,*F*2_). *e*_1_,*e*_2_, ⋯, *e*_6_ represent the residuals. Exploratory factor analysis estimates the phenotype-factor relationship from the data by allowing cross-loading. By choosing the largest factor loading value for each phenotype, phenotypes can be uniquely assigned to one of the two factors. In this example, Y1, Y2, and Y3 loaded on the F1 (with loadings of λ_11_, λ_21_, and λ_31_) and Y4, Y5, and Y6 loaded on F2 (with loadings of λ_42_, λ_52_, and λ_62_). Confirmatory factor analysis assumes that this relationship is known *a priori*.

### Multi-trait genomic best linear unbiased prediction

A Bayesian multi-trait genomic best linear unbiased prediction model was applied to partition inferred latent variables into genetic and environmental components.

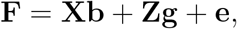

where **F** is the vector of estimated factor scores, **X** is the incidence matrix of covariates including the intercept and the top three principal components accounting for population structure, **b** is the vector of covariate effects, **Z** is the incidence matrix relating the factor scores of each latent variable to additive genetic effect, **g** is a vector of additive genetic effect, and **e** is the vector of residuals. Under the infinitesimal model of inheritance, **g** and **e** were assumed to follow a multivariate Gaussian distribution of **g** ~ *N*(0, ⊗_*g*_ ® **G**) and **e** ~ *N*(0, Σ_*e*_ ⊗ **I**), respectively. Here, **G** is a *n* × *n* genomic relationship matrix, **I** is a *n* × *n* identity matrix, ∑_*g*_ and Σ_*e*_ are variance-covariance matrices of additive genetic effect and residuals, respectively, and ⊗ is the Kronecker product. The **G** matrix was set as 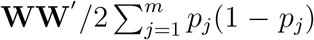, where **W** is the centered marker incidence matrix taking the values of 0 – 2_*p_j_*_ for zero copies of the reference allele, 1 – 2_*p_j_*_ for one copy of the reference allele, 2 – 2_p_*j*__ for two copies of the reference allele, and *p_j_* is the allele frequency at marker *j* = 1, ⋯, *m* (VanRaden, 2008). The prior distribution specifications followed those of Momen et al. (2019). A flat prior was assigned for **b**. The vectors of additive genetic and residual effects were assigned independent multivariate Gaussian priors with null mean and inverse Wishart distributions for the covariance matrices Σ_*g*_ and Σ_*e*_. A Gibbs sampler was used to obtain posterior distributions. A burn-in of 10,000 samples followed by an additional 90,000 samples, thinned by a factor of two, resulted in 45,000 available samples for posterior mean inferences. The MTM R package was used to fit the model (https://github.com/QuantGen/MTM).

### Bayesian network structure learning

The posterior means of genetic values of latent variables obtained from the Bayesian multi-trait genomic best linear unbiased prediction model were used to examine the manner in which the traits are interrelated using a Bayesian network. A Bayesian network is a graphical representation of the conditional independence among random variables based on a directed acyclic graph (Heckerman et al., 1995). For example, if an arrow arises from phenotype A to phenotype B, phenotype A is considered to impact phenotype B directly conditional on the remaining phenotypes, whereas the absence of an edge implies conditional independence given the remaining phenotypes. In this study, the Tabu search (Tabu) and Max-Min Hill-Climbing (MMHC) algorithms were applied to learn the underlying trait network structure of latent variables at the genetic level using the bnlearn R package (Scutari and Denis, 2014). These two algorithms were chosen because they yielded a reasonable result in a recent study (Yu et al., 2019). The Bayesian information criterion (BIC) score was calculated for the whole network and for each edge. A higher BIC score leads to greater model fit because the BIC score is rescaled by −2 in the bnlearn package. Additionally, the strength and uncertainty of the direction of each edge were estimated probabilistically by bootstrapping (Scutari and Denis, 2014). Before fitting the Bayesian network structure learning algorithms, genetic values of latent variables were transformed to be uncorrelated to meet the primary assumption of a Bayesian network (Töpner et al., 2017; Yu et al., 2019).

## Data availability

The data are available from the previously published studies. The agronomic, grain minerals, and rust related phenotypic data are available from Bhatta et al. (2018a,d, 2019) and the marker data are available from Bhatta et al. (2018d).

## Results

### Assessing factorability and factor selection

Figure 2 shows the Pearsons correlation coefficients among all observed variables represented in a heat map. Moderate to high correlations were observed within the spike-, mineral-, and rust-related traits. Because the objective of factor analysis is to model the interrelationships between observed traits with a smaller subset of latent variables, the presence of some block structures in the heat map suggests that our dataset is suited for factor analysis. This observation was supported by the overall Kaiser-Meyer-Olkin measure of sampling adequacy, which was estimated as 0.7, indicating that the factorability of the dataset was sufficient. Parallel analysis was performed to determine the appropriate number of latent variables. The first eight eigenvalues extracted from the original data were larger than the first eight eigenvalues obtained from simulated random data. Thus, eight underlying latent variables were examined in subsequent analysis.

**Figure 2:**
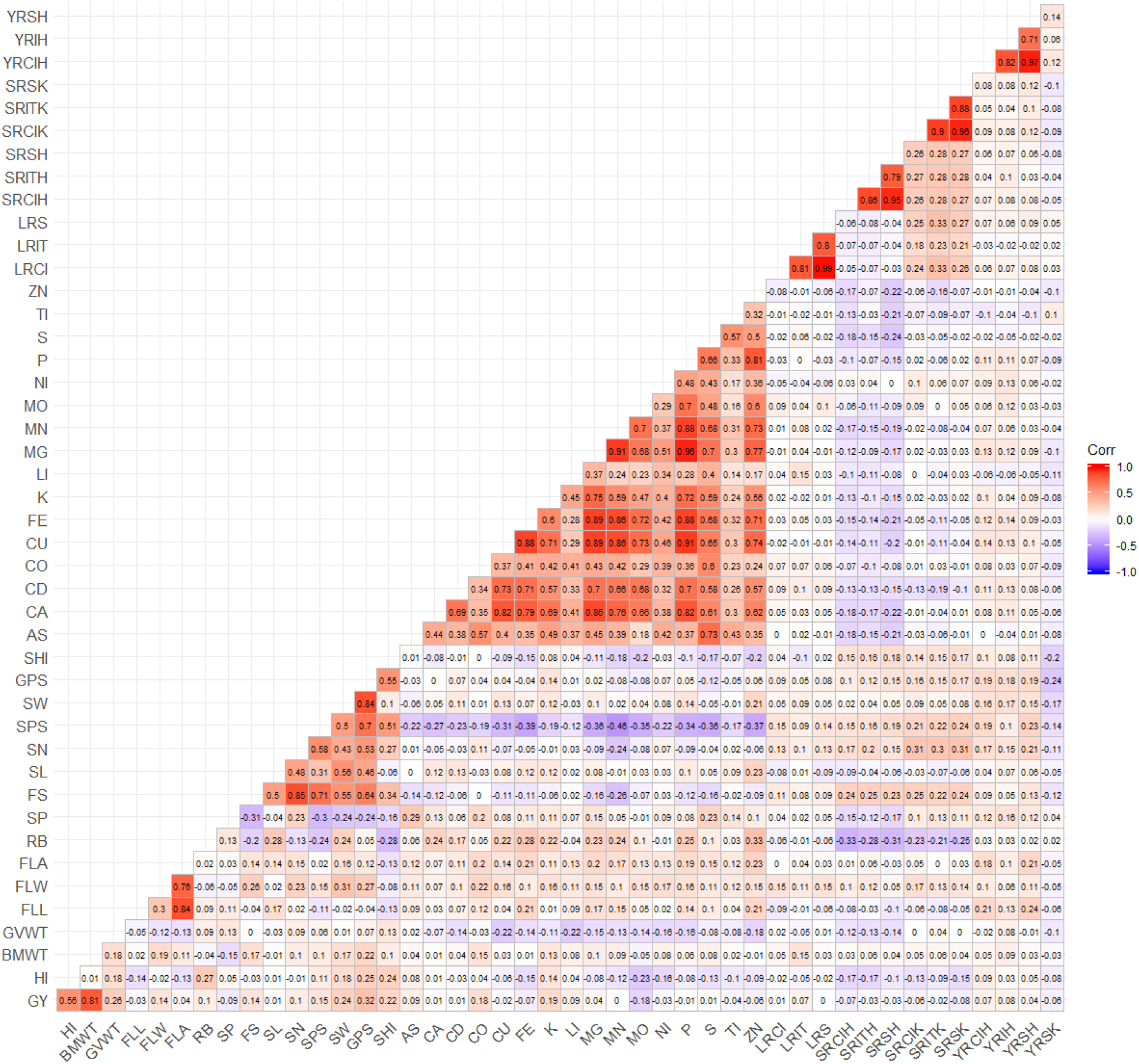
Pairwise Pearson’s correlations between 45 phenotypes. GY: grain yield, HI: harvest index, BWT: biomass weight, GVWT: grain volume weight, FLL: flag leaf length, FLW: flag leaf width, FLA: flag leaf area, SL: spike length, SN: spikelet number,SP: sterile spikelet, FS: fertile spikelet, RB: rachis break, SPS: seeds per spike, SW: spike weight, GPS: grain weight per spike, SHI: spike harvest index, AS: arsenic, CA: calcium, CD: cadmium, CO: colbalt, CU: copper, FE: iron, K: potassium, LI: lithium, MG: magnesium, MN: manganese, MO: molybdenum, NI: nickel, P: phosphorous, S: sulphur, TI: titanium, ZN: zinc, LRCI: leaf rust coefficient of infection, LRIT: leaf rust infection type, LRS: leaf rust severity, SRCIH: steam rust coefficient of infection at Haymana, SRITH: stem rust infection type at Haymana, SRSH: stem rust severity at Haymana, SRCIK: stem rust coefficient of infection at Kastamonu, SRITK: stem rust infection type at Kastamonu, SRSK: stem rust severity at Kastamanu, YRCIH: yellow rust coefficient of infection at Haymana, YRIH: yellow rust infection type at Haymana, YRSH: yellow rust severity at Haymana, YRSK: yellow rust severity at Kastamonu.

### Factor loading from EFA

Factor analysis was performed to understand the biological meaning of the eight latent factors by investigating the co-variation among measured observations using EFA. Figure 3 summarizes the degree of the contributions of unobserved factors to the observed phenotypes. Because EFA allows the cross-loading of phenotypes, an additional step is required so that each phenotype loads only on one factor. A heat map of the estimated factor loading values for each phenotype is shown in Figure 3A. The results showed that each variable had some nonzero loadings on several factors. Figure 3B shows the phenotype-latent variable pattern after selecting the largest loading for each phenotype and imposing a threshold of > |0.30|. This resulted in each phenotype loading on only one factor except for GVWT, RB, SP, and YRSK, which did not load on to any factors. The results showed that all mineral-related traits including As, Ca, Cd, Co, Cu, Fe, K, Li, Mg, Mn, Mo, Ni, P, S, Ti, and Zn were loaded on the first factor (F1) ranging from 0.34 to 0.98. Seven agronomic traits including FS, SL, SN, SPS, SW, GPS, and SHI were placed on the second factor (F2) and biologically all appear to be related to the plant structure. In this category, the lowest loading was estimated for the SHI (0.44) and the largest for GPS (0.91). The 12 disease-related phenotypes were distributed among 4 factors (F3, F4, F5, and F6) with a loading of at least 0.8 in their categories. FLL, FLW, and FLA traits with 0.84, 0.73, and 0.98 loadings, respectively, were placed on the seventh factor (F7). Finally, GY, HI, and BM loaded on the eighth factor (F8).

**Figure 3:**
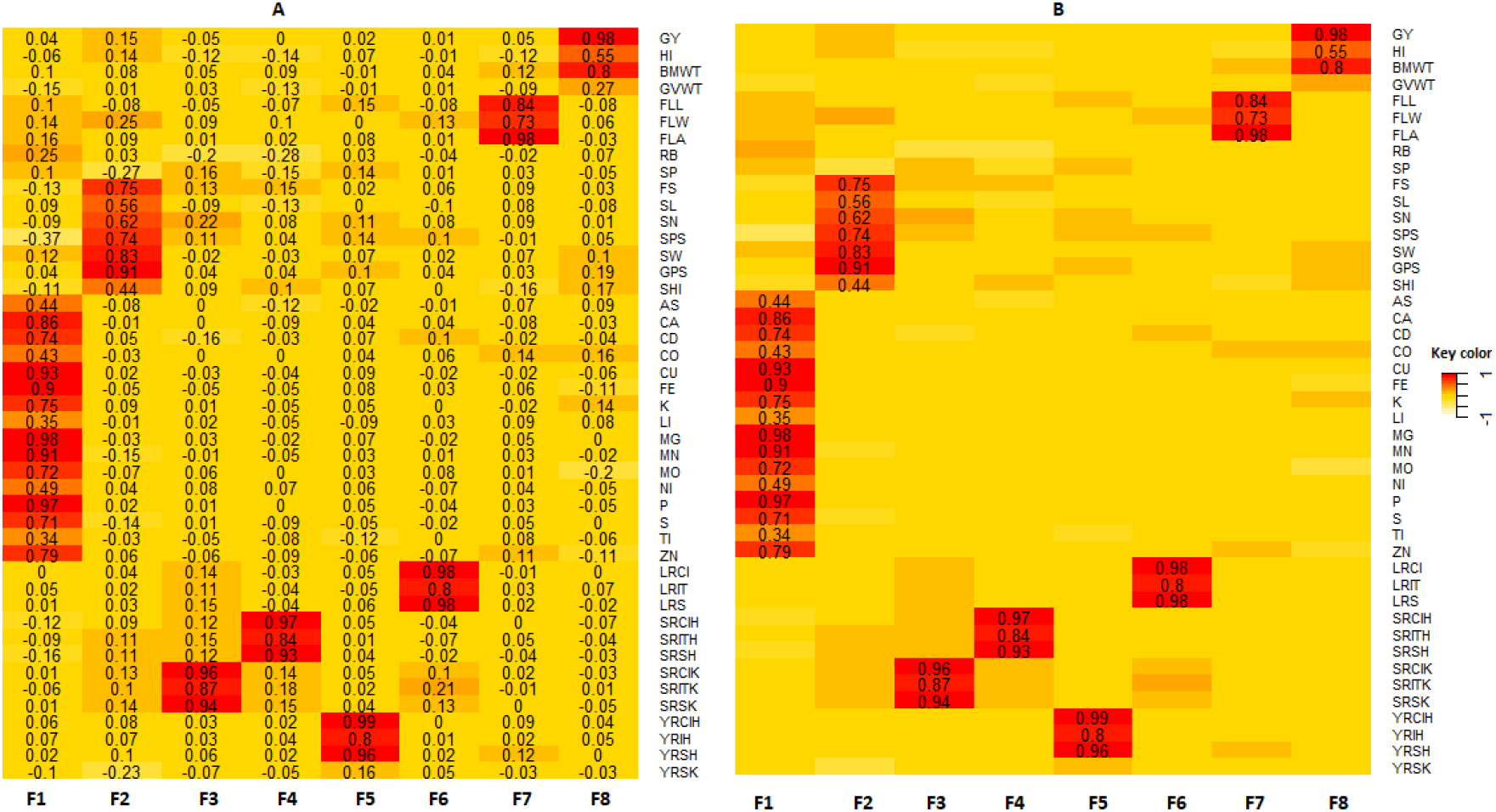
Panel A: heat map of factor loading values. Panel B: heat map of factor loading values after removing cross-loading by setting a cut-off value of |λ| > 0.30. The rows of each panel correspond to the observed phenotypes and the columns correspond to the eight factors (F1 to F8). Abbreviations of observed phenotypes are shown in Figure 2.

Figure 4 shows the overall inferred latent structure of the data. The biological meanings attached to the eight factors according to the EFA analysis were GYL: grain yield; ARC: plant architecture; FL: flag and leaf, MIN: minerals; YRD: yellow rust disease; SRDK: stem rust disease at Kastamonu; SRDH: stem rust disease at Haymana; and LRD: leaf rust disease. These estimated latent factors were subsequently evaluated to determine their genetic interrelationships.

**Figure 4:**
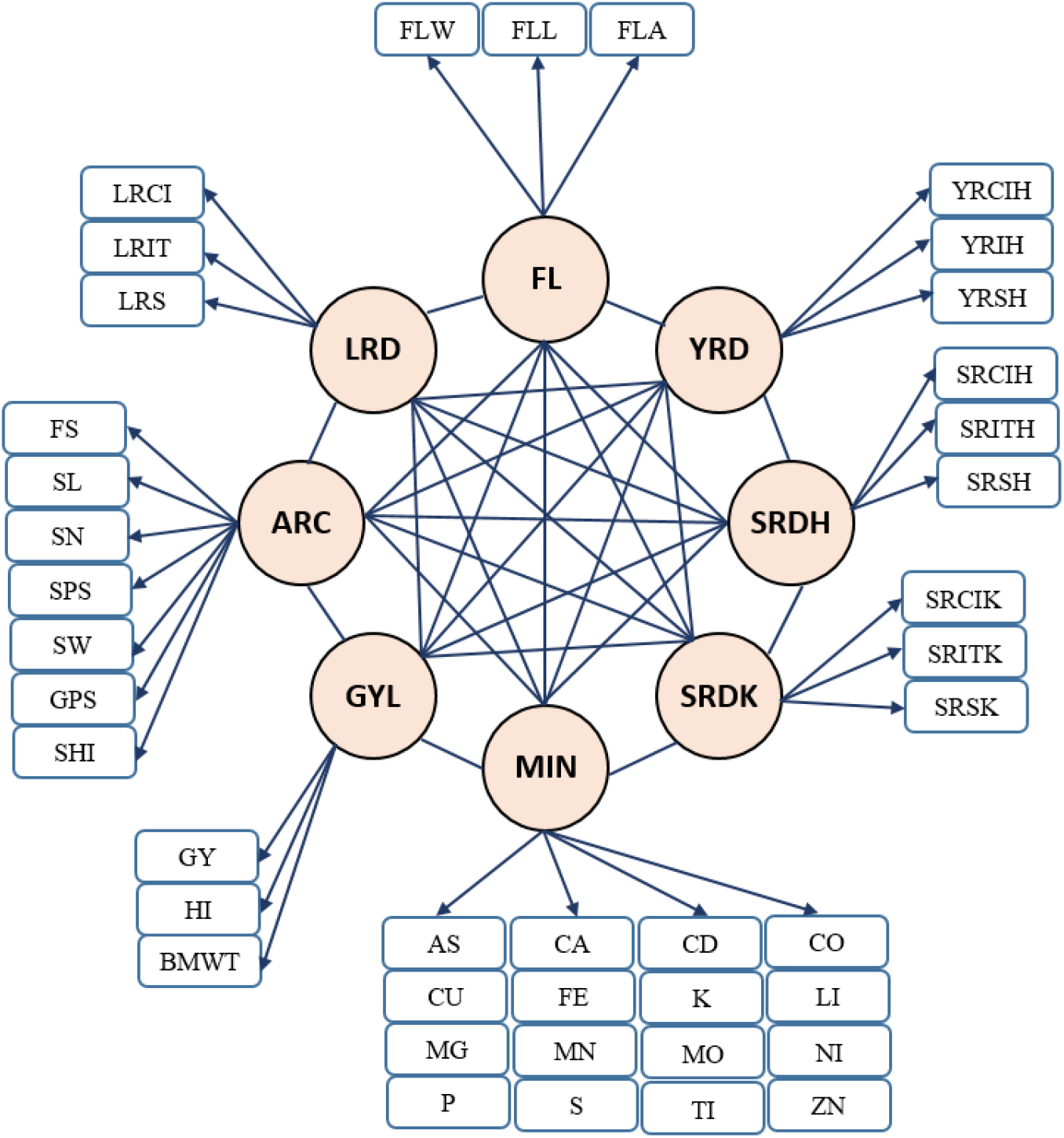
Relationship between eight latent variables and observed phenotypes based on exploratory factor analysis. GYL: grain yield related traits, ARC: architecture related trait, FL: flag and leaf related traits, MIN: mineral-related traits, YRD: yellow rust related traits, SRDK: stem rust related traits at Kastamonu, SRDH: stem rust related traits at Haymana, LRD: leaf rust related traits. The eight latent factors were assumed to be correlated. Abbreviations of observed phenotypes are shown in Figure 2.

### Confirmatory factor analysis

Table 1 shows the posterior means and their posterior standard deviations of the standardized loadings, PSRF, and R^2^ statistics from the Bayesian CFA. Convergence was diagnosed from the PSRF of each observed phenotype. The estimated PSRF values for all phenotypes were close to 1, suggesting that they converged to a stationary status. The result showed that the eight latent factors strongly contributed to the observed phenotypes. For the latent factor GYL, the lowest and highest loading values were obtained for HI and GY, respectively. For the FL latent factor, all three phenotypes presented a loading of at least 0.77. In ARC, the factor loading values varied from SHI to FS in ascending order. The MIN latent factor was associated with the 16 observed phenotypes, which was the largest factor. The lowest and highest loading values were obtained for Ti and Mg, respectively. The remaining four latent factors including LRD, SRHD, SRKD, and YRD, which are relevant to diseases, showed that the data fit well with >0.8 loading. The extent of R^2^ values mostly agreed with the estimated loadings with a correlation of 0.99.

**Table 1:**
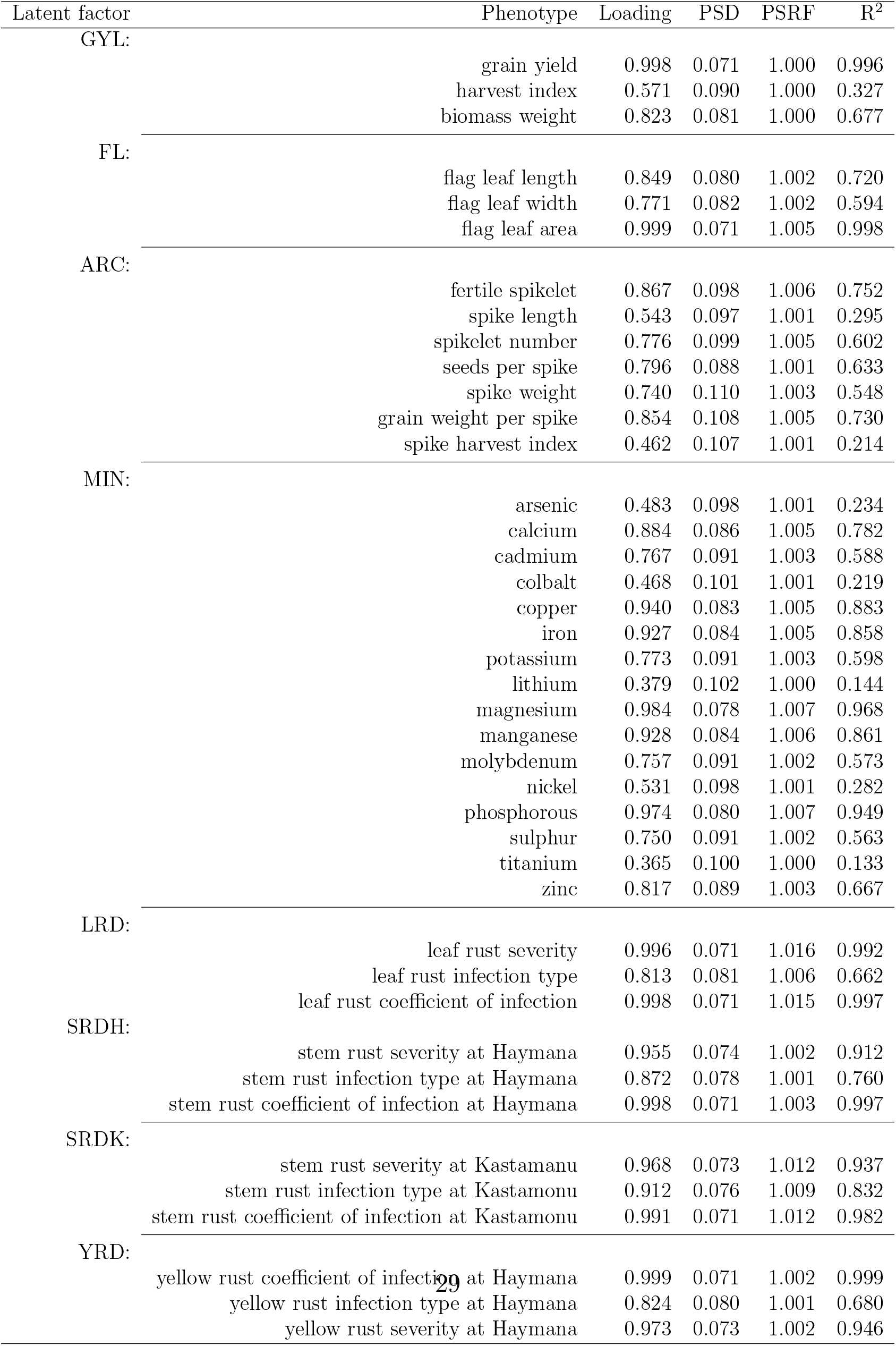
Factor loading values from the Bayesian confirmatory factor analysis. PSD: posterior standard deviation, PSRF: potential scale reduction factor, GYL: grain yield, ARC: plant architecture, FL: flag and leaf, MIN: mineral-related traits, YRD: yellow rust, SRDK: stem rust at Kastamonu, SRDH: stem rust at Haymana, LRD: leaf rust, and R^2^: coefficient of determination.

### Bayesian network among genomic latent factors

The Bayesian network was used to investigate the interrelationships among the genetic components of latent factors. Because SRDH and SRDK capture the same set of phenotypes with a high correlation (Figure 3) but were collected at different locations, only SRDH was used for trait network structure analysis. As shown in Figure 5, Tabu yielded six directed edges from FL to LRD and MIN, from YRD to LRD and GYL, from MIN to ARC, and from SRDH to GYL. However, MMHC only produced three directed edges that were a subset of the Tabu network. Thus, the consensus network has common directed edges from FL and LRD, from YRD to GYL, and SRDH to GYL. These results suggest that there is stronger evidence that FL, YRD, and SRDH directly influence LRD, GYL, and GYL, respectively. In both networks, the bootstrapping results revealed that confidence was always higher regarding the presence or absence of edges compared to the directions of edges. The goodness-of-fit statistics measured by BIC is shown in Table 2. This table shows how well the paths mirror the dependence structure of the data. According to the BIC values, Tabu yielded a larger BIC score than the MMHC algorithms for the entire network (−423.61 vs. −437.39). For each specific path, removing SRDH → GYL resulted in the largest decrease in the BIC score, suggesting that this path plays the most important role in the network structure. This was followed by YRD → GYL and FL → LRD. The top three most influential paths in Tabu formed the network structure of MMHC.

**Figure 5:**
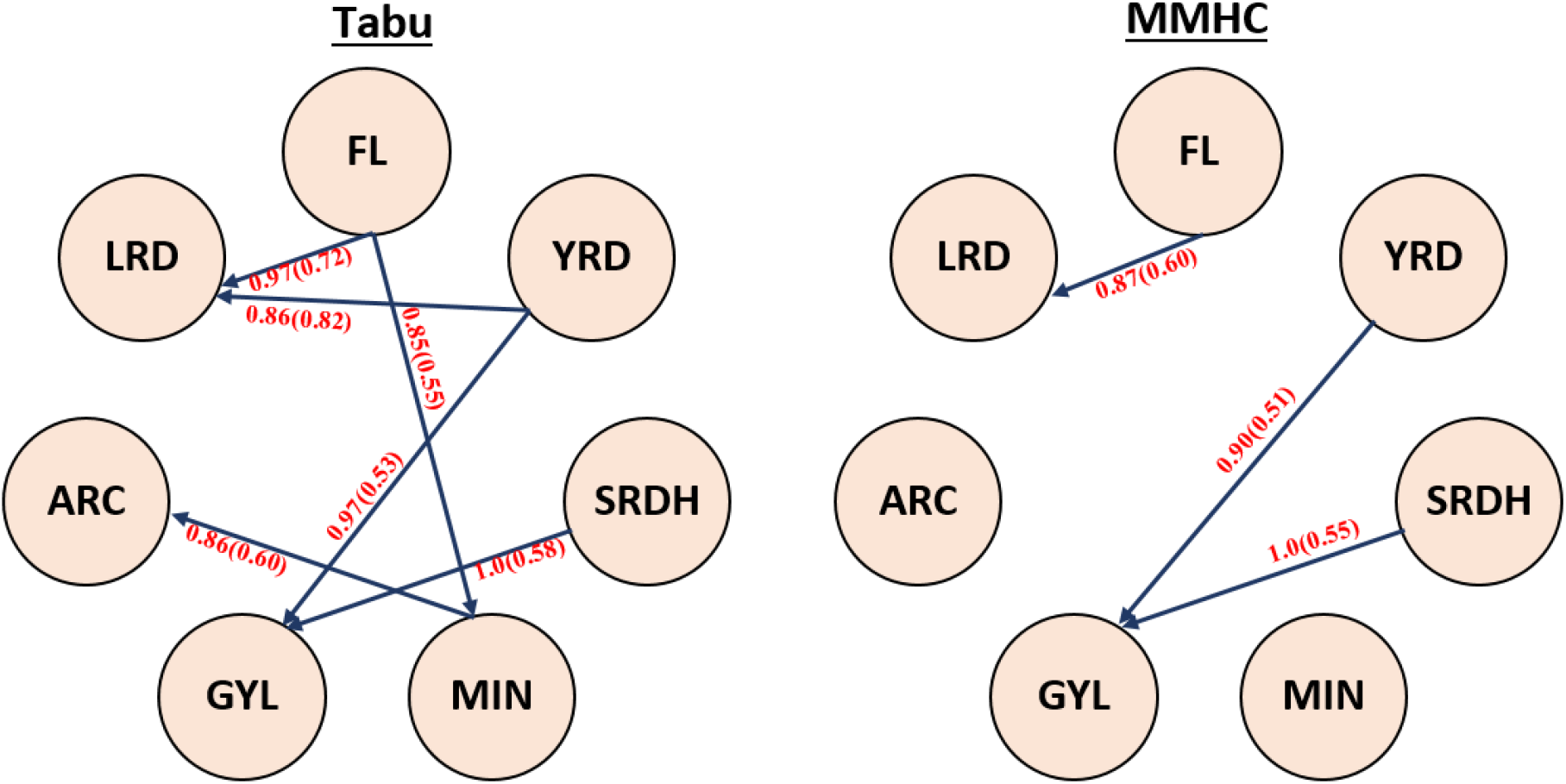
Bayesian networks learned from Tabu search (Tabu) and Max-Min Hill-Climbing (MMHC). Structure learning test was performed with 5,000 bootstrap samples. Labels of the edges refer to the strength and direction (parenthesis) which measure the confidence of the directed edge. The strength indicates the frequency of the edge is present and the direction measures the frequency of the direction conditioned on the presence of edge. GYL: grain yield related traits, ARC: architecture related trait, FL: flag and leaf related traits, MIN: mineral-related traits, YRD: yellow rust related traits, SRDK: stem rust related traits at Kastamonu, SRDH: stem rust related traits at Haymana, LRD: leaf rust related traits.

**Table 2:** Bayesian information criterion (BIC) scores for pairs of nodes reporting the change in the score caused by an arc removal relative to the entire network score. Tabu: Tabu Search, MMHC: Max-Min Hill-Climbing, GYL: grain yield traits, FL: flag and leaf traits, MIN: mineral traits, ARC: architecture traits, LRD: leaf rust disease, SRDH: steam rust disease at Haymana, and YRD: yellow rust disease.

| Algorithm | from | to | BIC |
| --- | --- | --- | --- |
| Tabu | FL | MIN | -2.074 |
|  | FL | LRD | -9.648 |
|  | MIN | ARC | -5.884 |
|  | SRDH | GYL | -32.297 |
|  | YRD | GYL | -16.399 |
|  | YRD | ARC | -5.916 |
| MMHC | FL | LRD | -5.989 |
|  | SRDH | GYL | -32.297 |
|  | YRD | GYL | -16.3997 |

## Discussion

### Data-driven latent variable analysis

With the availability of large volumes of measured observations per individual because of recent advances in phenomics, it is critical to develop a phenotype-centric statistical approach. Factor analysis is an effective method for handling many response variables in a quantitative genetic framework (Runcie and Mukherjee, 2013; Peñagaricano et al., 2015; Rocha et al., 2018; Yu et al., 2019, 2020). The central idea behind factor analysis is to model the observed phenotypes through unobserved latent factors by maximizing the common variance between correlated phenotypes. In the current study, latent factors were directly inferred from the field data of physiological and morphological phenotypes in wheat using EFA followed by estimating their factor scores by CFA. This allowed the analysis of the lower dimensional data because the number of latent factors was less than the number of observed phenotypes. The combination of EFA and CFA enabled the evaluation of the genetics of latent factors that were predicted to give rise to the observed phenotypes. Our results demonstrate that a data-driven approach for estimating latent factors using EFA is useful because the observed traits were uniquely assigned to one of the factors with biological interpretations. This contrasts with the results of a recent study by Yu et al. (2019), in which observed phenotypes were classified into factors based on prior biological knowledge. However, in most scenarios, the phenotype-latent variable pattern may be unknown. In contrast, EFA can be used to perform latent variable analysis by estimating latent factors from data when the latent structure cannot be determined *a priori*.

The interrelationships among latent variables were investigated at the genomic level using Tabu and MMHC. Based on the BIC values, Tabu resulted in a better fit than MMHC. This agrees with the findings of recent studies using Bayesian networks (Töpner et al., 2017; Scutari et al., 2018; Yu et al., 2019). The trait network structure inferred from MMHC was a subset of that of MMHC. Additionally, the three directed paths identified from MMHC were the top three most important paths in Tabu according to BIC. This suggests that the networks structures were consistent between Tabu and MMHC. Thus, the trait network derived from MMHC can be considered the consensus network that is more reliable. The network structures from Tabu and MMHC may become aligned by increasing the sample size. Inferring a trait network from observational data is an emerging topic in quantitative genetics (Valente et al., 2010). Because breeders are often interested in the impact of external intervention or the selection of one trait over other traits, distinguishing undirected edges from directed edges is important. The trait network learned in this study can also be integrated into SEM-GWAS, which is a framework to perform multi-trait genome-wide association analysis derived from structural equation models (Momen et al., 2018, 2019). The combination of data-driven EFA and Bayesian network approaches is particularly useful for analyzing image-based high-throughput phenotyping data, where relationships within image-based phenotypes and between classical phenotypes and image-based phenotypes may not always be obvious.

### Biological meaning of the inferred relationships

Previous studies revealed the negative genetic associations of yellow and stem rust traits with grain yield traits. Wheat rust diseases are foliar fungal diseases whose infection on the flag leaf close to the grain filling period causes a decline in the photosynthetic ability of the plant, drastically decreasing the grain filling process and reducing the biomass yield, thousand kernel weight, and harvest index (He et al., 2019; Bhatta et al., 2018a; Herrera-Foessel et al., 2006). Thus, the reduction of these important traits results in a reduction in the final grain yield (SRDH → GYL and YRD → GYL). Wheat leaf rust may be affected by flag leaf traits such as FLL, FLW, and FLA (FL → LRD). As the flag leaf area increases, the surface also becomes greater, increasing the risk of disease infection on the wider and longer leaves.

Flag leaf traits play important roles in the synthesis, translocation, and remobilization of photo-assimilates and minerals to the grains (Sperotto et al., 2013). A recent study on *Triticum sps.* showed that the flag leaf contains two-to three-fold higher concentrations of Fe and Zn than the grain mineral concentrations (Hu et al., 2017). They also found strong positive correlations between leaf and grain Fe and Zn concentrations. Another study used more than 120 hexaploid wheat lines and reported a significant positive correlation of flag leaf N concentrations at anthesis with grain Fe, Mn, and Cu (SHI et al., 2013). These results suggest that flag leaf traits play an important role in determining the grain mineral concentration, which agrees with our results indicating a direct link from FL to MN.

Foliar diseases such as yellow rust, caused by *Puccinia striiformis f. sp. tritici (Pst)*, is an important foliar fungal disease of wheat that causes major yield loss (Bhatta et al., 2019). This disease produces rust pustules on leaves and reduces the process of photosynthesis and translocation of photosynthate to grain yield traits, which in turn inhibit grain filling, possibly resulting in a significant reduction in grain weight and ultimately reducing grain yield (Ye et al., 2019; Murray and Murray, 2005). A recent study on winter wheat germplasm showed that yellow rust infection seriously damaged the photosynthetic function of leaves at an earlier stage of grain filling, leading to biomass loss (He et al., 2019). Additionally, the presence of foliar diseases in wheat is associated with a reduction in the biomass weight and harvest index by reducing the healthy leaf area and affecting healthy spike growth (Gooding et al., 2000; Dimmock and Gooding, 2002), indicating that yellow rust traits affected grain yield-related traits (YRD → GYL).

Several studies have reported negative associations between grain minerals and architecture-related traits. A larger number of seeds per spike and kernel size in wheat is associated with lower grain mineral accumulation in the grain, which is mainly attributed to the grain mineral dilution effect (Bhatta et al., 2018a; Guttieri et al., 2015). Similarly, the nitrogen concentration in the grains depends on their position within the spike Calderini and Ortiz-Monasterio (2003); Herzog and Stamp (1983), suggesting that spike architecture traits have important impacts on grain mineral traits (MIN → ARC).

## Conclusions

This study demonstrates that data-driven latent variable analysis can reveal the underlying structure of phenotypes on a smaller dimensional scale. Thus, determining the genetic effects of correlated traits by factor analysis is an efficient approach for learning the minimum set of core factors contributing to high-dimensional observed phenotypes. Additionally, by reconstructing a more general structure of genomic latent factors from observed phenotypes using a Bayesian network, a clearer picture of trait interdependency can be obtained, which is useful for developing breeding and management strategies for crops such as wheat.

## Acknowledgements

This work was supported in part by Virginia Polytechnic Institute and State University startup funds to GM.

## Author contributions

MM, MB, WH, and GM conceived the study. MM and HY analyzed the data. MM drafted the manuscript. WH, MB, HY, and GM revised the manuscript. GM supervised and directed the study. All authors read and approved the manuscript.

## Conflict of interests

The authors declare that they have no competing interests.

## Notes

### Competing Interest Statement

The authors have declared no competing interest.

### Summary of Updates

There were typos in the title and abstract. explanatory -> exploratory.

